# DynaRepo: The repository of macromolecular conformational dynamics

**DOI:** 10.1101/2025.08.14.670260

**Authors:** Omid Mokhtari, Emmanuelle Bignon, Hamed Khakzad, Yasaman Karami

## Abstract

Proteins, RNA, and DNA are central to virtually all cellular processes, often assembling into macro-molecular complexes to perform their functions. While these molecules are inherently dynamic, most methods for studying their mechanisms focus on static structures. Recent deep learning advances in protein structure prediction highlight the potential of data-driven approaches, yet dynamic behavior that is critical for interactions such as antibody–antigen recognition, intrinsically disordered proteins, and protein–nucleic acid binding, remains underexplored. To address this gap, we present DynaRepo, a repository of macromolecular conformational dynamics comprising ~450 complexes and ~270 single-chain proteins from PDBbind, the Structural Antibody Database (SAbDab), and benchmark sets. Each complex was simulated in triplicate for 500 ns, totaling >1100 *µ*s of molecular dynamics data, with extensive pre-calculated analyses. DynaRepo provides a foundation for dynamics-aware deep learning frameworks and is freely available at: https://dynarepo.inria.fr/.

## Introduction

Biological molecules often interact together and form complexes to execute their functions. These functions are directly linked to the arrangement of atoms in 3D space (structure) and to their motions (dynamics). Macromolecular complexes undergo conformational changes in response to different environmental conditions, such as mutations, changes in temperature or electrostatic potential, as well as binding to other molecules. These changes may alter the structural plasticity, induce malfunctioning, and thereby provoke diseases. Therefore, characterizing the structure and conformational dynamics of macromolecular complexes can help to understand the molecular mechanisms of underlying diseases and ultimately to design drugs that modulate these mechanisms.

With the transformative advances brought by deep learning–based methods such as AlphaFold2 [1], AlphaFold3 [2], and RoseTTAFoldNA [3], the next major challenge is to develop dynamics-aware approaches. This need is critical, as the conformational heterogeneity of macromolecules underpins many biological functions that current methods cannot capture. Yet, predicting macromolecular dynamics remains difficult, largely due to the scarcity of openly available, high-quality dynamic datasets. Molecular dynamics (MD) simulations offer a powerful and versatile computational microscope to address this gap, generating conformational ensembles that reveal molecular motions and capture the dynamic heterogeneity essential for function.

Over the past few decades, MD simulations have become crucial for elucidating the conformational behavior of proteins. Short trajectories on the order of tens of nanoseconds have proven invaluable for generating protein structure ensembles, thereby improving docking performance, pinpointing interaction sites, and revealing flexibility patterns that identify residues participating in protein–protein interfaces. Simulations extending to hundreds of nanoseconds have enabled the mapping of allosteric pathways, while even longer trajectories have shed light on large-scale conformational rearrangements. However, comparative analyses across simulations from different research groups remain challenging due to variations in system preparation, MD protocols, software packages, and force fields. To address these discrepancies, several initiatives have emerged to create shared repositories for MD data, such as MDRepo (open repository for MD simulations of proteins) [4] and MDDB (a federated system of topical simulation repositories). Other resources focus on specific protein classes, including MemProtMD for membrane proteins [5], GPCRmd for G protein–coupled receptors [6], and SCoV2-MD for SARS-CoV-2 proteins [7]. More general-purpose databases, such as MoDEL [8], Dynameomics [9], and ATLAS [10], provide MD trajectories for a broad range of soluble proteins.

MDDB is a European initiative to establish the first unified, distributed database for MD simulations, enabling scientists to share, access, and build upon one another’s work https://mddbr.eu/. Conceptually, it operates as a federated network in which multiple nodes are connected to a central core hosted in Barcelona. MDDB provides standardized infrastructure and protocols for all nodes (referred to as MD Posits), while the MD data itself is generated by researchers both inside and outside the consortium. Each collaborator manages their own node, which is fully integrated with the main MDDB server.

In this context, we have developed DynaRepo, the first MDDB node established in France, dedicated to large-scale simulations of macromolecular dynamics. Unlike most existing repositories, DynaRepo focuses on macromolecular complexes including protein–nucleic acid assemblies rather than isolated proteins or small molecules. By incorporating these data into a structured MDDB node, DynaRepo complements and extends current resources. The repository’s time-resolved trajectories explicitly capture conformational flexibility, providing a rich foundation for training AI models in binding site prediction, functional annotation, stability assessment, and binding affinity estimation. These capabilities are crucial for biology, biophysics, and therapeutic discovery.

## MATERIALS AND METHODS

### Data selection

DynaRepo consists of MD simulations on entries from four distinct sets: protein-protein complexes from PDBbind version 2020 [11], a set of transient proteins, antigens selected from the structural antibody database (SAbDab) [12], and a set of nucleosome complexes. The distribution of entry sizes based on number of residues is shown for each dataset in **Supplementary Fig. S1**.

#### Protein-protein complexes from PDBbind

We selected protein–protein complexes from the PDB-bind dataset (2,852 entries) using the following filtering pipeline: *i)* Complexes determined by NMR (200 entries) were excluded. *ii)* Only complexes with a resolution better than 3.5 Åwere retained, leaving 2,526 entries. *iii)* Structural gaps were identified using pdb-tools [13]; only structures with fewer than four gaps were kept, reducing the dataset to 1,185 entries. *iv)* All-against-all structural alignments were performed using MM-align [14]. Pairwise distances were computed as 1 − MM-score, followed by hierarchical agglomerative clustering (average linkage). Clusters were extracted using a distance threshold of 0.55, yielding 433 clusters. *v)* After manual inspection, 405 representative structures were chosen for subsequent simulations.

#### Transient benchmark

A set of 53 single-chain proteins reported to be involved in highly transient interactions [15] was selected. Overlapping entries with those selected from PDBbind (previous paragraph) were removed. We performed MD simulations on both the individual chains and the corresponding complexes.

#### Antigens

The antigen set was made initially using 105 unique and diverse single-chain antigens provided by Zhao et al. [16]. To further extend this dataset, we utilized the SAbDab database [12], keeping only protein-type antigens derived from X-ray crystallography with a resolution better than 3.0Å. We further filtered for entries with both heavy and light chain antibodies and available reported affinity data. Gap information, including the number and lengths of gaps, was extracted using pdb-tools [13]. Sequence-based clustering was performed using mmseqs2 [17] with a similarity threshold of 0.3, resulting in 174 clusters. After manual inspection, 116 antigen chains were selected for further MD simulations.

#### Nucleosomes

A set of simulations of the nucleosome core particle, a complex of eight histone proteins around which 145-147 base pairs of DNA are wrapped, was included from previous studies. MD data of the canonical nucleosome core particle with either the *α*-satellite (PDB ID 1KX5, 6 replicas of 2 *µ*s) [18] or the 601 Widom sequence (3LZ0, 3 replicas of 4 *µ*s) [19] were added, as well as simulations with different cysteine oxidative post-translational modifications mutated from the 1KX5 structure (6 replicas of 2 *µ*s with single or double S-sulfonylated H3C110 [18], and 5 replicas of 2 *µ*s with either S-sulfonylated H3C110 or S-nitrosylated H3C110 [20]). Finally, 2 replicas of 4*µ*s of the nucleosome core particle bound to the SIRT6 deacetylase were also included (PDB ID 8OF4) [19].

### Molecular Dynamics Simulations

#### Protocol for proteins

For every system, we performed three replicates of 500 ns MD simulations starting from the experimental structure. First we removed all the ligands, but kept the ions. Then, missing residues were added using MODELLER for consecutive gaps of maximum 5 residues or with AlphFold3 if the gap was longer than 5 amino acids. For the AlphaFold3 predictions, we chose the one with the highest pLDDT score. MD simulations were carried out with the GROMACS 2024.2 [21] using the CHARMM36m force field parameter set: (i) Na^+^, Cl^−^ counter-ions were added to reproduce physiological salt concentration (150 mM solution of sodium chloride), (ii) the solute was hydrated with a dodecahedron box of explicit TIP3P water molecules with a buffering distance of up to 12 Å, and (iii) hydrogen atoms were added and the environment of the histidine was checked using the Reduce software [22]. For each system, the energy minimization was performed by steepest descent algorithm for 5000 steps, to minimize any steric overlap between system components. This was followed by an equilibration simulation in an NPT ensemble at 310 K, allowing the lipid and solvent components to relax around the restrained protein. All the protein non-hydrogen atoms were harmonically restrained, with the constraints gradually reduced in 6 distinct steps with a total of 0.375 ns. During the system equilibration steps, the pressure was maintained at 1 bar with Berendsen thermostat and barostat [23], respectively. For every system, three replicates of 500 ns, with different initial velocities were performed in the NPT ensemble using the time step of 2.0 fs. The temperature was kept constant at 310 K using V-rescale thermostat [24] and a constant pressure of 1 atm was maintained with C-rescale barostat [25]. Isotropic pressure treatment was applied to the system. The bonds involving hydrogen atoms were constrained using the LINCS algorithm [26], while the electrostatic interactions were calculated using Particle Mesh Ewald method [27], and the coordinates of the system were written every 100 ps. **Supplementary Fig. S2** highlights the overall evolution of per-residue fluctuations and **Supplementary Fig. S3** shows the RMSD values along the simulation for all the entries in the database. RMSD values are reported with reference to both the initial structure and the average structure (in blue and purple, respectively).

#### Protocol for protein-nucleic acid complexes

All simulations of protein-nucleic acid complexes were taken from previous works [18–20], performed with either Amber20 [28] or NAMD3 [29]. The nucleosome systems were modeled using a combination of the ff14SB [30], BSC1 [31], CUFIX [32], and TIP3P [33] force field parameters. Each system was soaked in a truncated octahedral water box built with a 20 Å buffer, and NaCl counterions were added to achieve a 150 mM ionic strength. A 10,000 to 30,000 steps minimization was carried out, followed by a 40 to 100 ns NPT equilibration with decreasing restraints on the solute atoms. All simulations were performed at a 300 K temperature (Langevin thermostat [34]) and a 1 atm pressure (Langevin piston [35] or Berendsen thermostat [23]). A 4.0 fs time step was employed, allowed by the use of the SHAKE [36] and Hydrogen Mass Repartitioning algorithms [37]. Electrostatic interactions were treated using the Particle Mesh Ewald approach [27]. Production runs were performed in 2 to 6 replicas spanning from 2 to 4 *µ*s, on systems accounting for 300,000 to 400,000 atoms. Therefore, simulations were stripped before deposition on DynaRepo to lighten the trajectory files by setting 1 frame = 1 ns.

### Database implementation

DynaRepo follows the standard workflow of MDDB for data analysis, metadata preparation, and visualization using MDPosit and a REST API interface [38]. The workflow is available in GitLab: https://mmb.irbbarcelona.org/gitlab/d.beltran.anadon/MoDEL-workflow. It integrates a suite of biomolecular analysis tools into a reproducible pipeline that supports major MD engines such as AMBER, GRO-MACS, NAMD, OpenMM, and Desmond. This workflow performs both standard and system-specific analyses, which are automatically uploaded to the DynaRepo database along with trajectory, topology, and rich metadata. The setup includes automated enrichment via external biological databases and supports programmatic access through a REST API, enabling advanced, customizable queries and data retrieval. To promote interoperability, an OPTIMADE-compliant interface is also provided, bridging life sciences and materials research standards. The DynaRepo node is hosted and maintained by the Inria Center at the University of Lorraine.

DynaRepo accepts contributions from other users, which can be made by contacting the authors and uploading the MD data. With support from the Inria Center, we then perform the automatic post-simulation analysis to generate the metadata and upload all results to the DynaRepo server.

### Post-simulation analysis

A comprehensive set of post-simulation analyses is pre-computed via the MDDB workflow and provided for each entry. The available analyses are listed below (technical details are available on the help page).

**Root mean squared deviation (RMSD)** is computed against the first and the average structure of the trajectory, with possibility to select different atoms (all atoms, heavy atoms, and backbone and alpha carbon). RMSD is also calculated per residue (first frame of each residue and its conformations in up to 200 frames), and pairwise (for interface residues in complexes).

**TM score** are calculated against the first frame and the average structure of the trajectory [39].

**Radius of gyration** are computed about the x-, y- and z-axes, as a function of time.

**Fluctuation** analysis computes the root mean square fluctuation (RMSF) of atomic positions along the simulation time after aligning to the first frame.

**Principal component analysis (PCA)** calculates the covariance matrix (up to 2000 frames) along the simulation time (only with backbone atoms) in order to obtain a set of eigenvectors and its associated eigenvalues.

**Solvent accessible surface (SAS)** analysis computes the area of contact between the solvent and each residue along simulation time.

**Clusters** calculation allows partitioning of the trajectories. The user can adjust both the number of clusters and the number of links. It also shows clusters based on different set of interacting chains.

**Distance per residue** analysis computes distance (average and standard deviation) between each pair of residues in each interaction interface over the simulation time.

**Electrostatic potential surface** illustrates interface residues (in complexes) in surface and in balls+sticks modes.

**Hydrogen bonds** are considered when the distance between an acceptor and a donor atom is lower than 3 Å and the ‘acceptor - hydrogen - donor’ atoms angle is higher than 135 degrees.

**Energies** analysis computes the Electrostatic and Van der Waals forces for all residues along the simulation time using CMIP [40]. Energy averages per residue are calculated for the overall, start and end of the trajectory. Energies are calculated for each atom individually and then added by residues.

**Pockets** analysis is calculated by MDpocket [41] which searches for cavities over the simulation time.

### Computational complexity of the workflow

The computational time required for the workflow depends strongly on the number of chains and residues. For single-chain trajectories, the calculations typically take from a few minutes up to a few hours per replica, depending on the system size. However, the runtime increases drastically when more than one chain is present in the studied system. In the case of nucleosome simulations that are the largest systems in the database, the computational time for the analysis workflow reached about 60 hours per replica. This significant increase is mainly due to the additional calculations of hydrogen bonds, per-residue distances, electrostatic potential surfaces, energies, and pockets. These analyses are performed only between interacting chains, not within a single chain.

## RESULTS

The DynaRepo database is a growing collaborative initiative designed to host MD simulations generated both in-house and by external contributors. Built to comply with FAIR principles (Findable, Accessible, Interoperable, and Reusable), DynaRepo facilitates open science by enabling the reuse and exploration of MD trajectories by the broader research community [42]. As an independent node within the federated MDDB system, DynaRepo contributes to a global effort to provide accessible data on macromolecular conformational dynamics.

Currently, the database contains simulations for 405 macromolecular complexes, carefully selected and clustered based on sequence and structural similarity from the PDBbind dataset; 220 single-chain antigens sourced from the SAbDab database; and the “Testing Transient” dataset, which includes 53 protein–protein complexes in both bound and unbound forms. Each complex has been simulated in three replicates of 500 ns, yielding a cumulative simulation time of approximately 1065 *µ*s and encompassing around 1500 protein chains. Additionally, simulations of seven structural variants of the nucleosome, a protein–DNA complex involved in chromatin organization, are available, contributing another 81 *µ*s of MD data.

Each entry in DynaRepo is enriched with extensive post-simulation analyses, accessible through interactive visualizations. These include RMSD, RMSF, TM-score, radius of gyration, principal component analysis (PCA), solvent-accessible surface area, per-residue distance metrics, electrostatic potential surfaces, hydrogen bonding patterns, energy profiles, and pocket detection.

### Overview and trajectory pages

Every entry has a unique accession number, a name, a small preview of the 3D structure and the list of available analyses (**Fig. 1A**). The name of the entry shows database name (DynaRepo) and the PDB code. After selecting an entry from the BROWSE page, the user can select between different replicas of the simulation. For each replica there are four tabs: OVERVIEW, TRAJECTORY, ANALYSES and DOWNLOAD. The OVERVIEW page provides a summary of the simulation protocol, information on the structure (the sequence, PDB code, UniProt ID, etc.) and citations. The TRAJECTORY page plays the movie from the MD simulation trajectory, where you can select either of the partners (proteins or NA) in the complex to see their behavior during the simulation time (**Fig. 1B**). Below the visualization panel, functional analysis can be seen (**Fig. 1C**), where the functional annotation for each chain in the complex is represented. These functional annotations including functional families (FunFam), domains, and sites are mapped over sequence and derived from external resources such as InterPro [43], and GENE3D [44].

**Figure 1:**
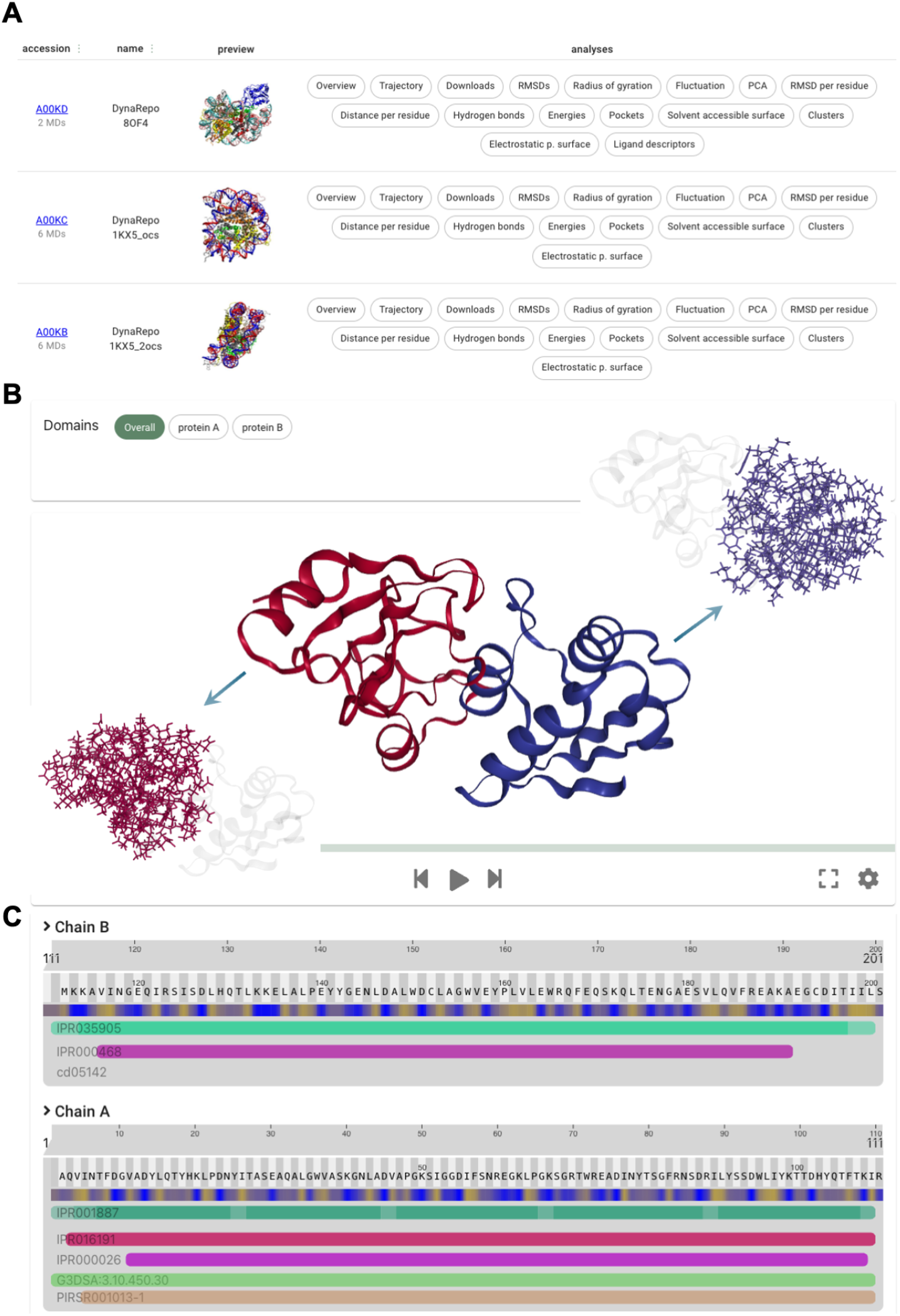
An overview of the DynaRepo database. **A)** For each entry there are: a unique accession number, a name, a 3D structure and the list of all the analyses. **B)** The Trajectory page plays the trajectory along the time, and **C)** offers functional analysis mapped over the sequence.

### Macromolecular analyses page

A large number of analyses are pre-calculated for each entry. A selection of such analyses are shown in **Fig. 2** for one entry of the database. These analyses are detailed in the Methods section and two representative examples are shown in the Supplementary data (**Supplementary Fig. S4-15**). The analyses grouped in three broad categories of: Quality control, Interactions, and Other. The quality control features include RMSDs (per residue, and pairwise) **Fig. 2A**, solvent accessible surface **Fig. 2B**, fluctuation **Fig. 2C**, radius of gyration **Fig. 2D**, PCA, and clusters **Fig. 2E**. Interactions contain information regarding the distance per residue, electrostatic potential surface **Fig. 2F**, hydrogen bonds, and energies. And the Other features represent pockets **Fig. 2G**, ligand descriptors, and density.

**Figure 2:**
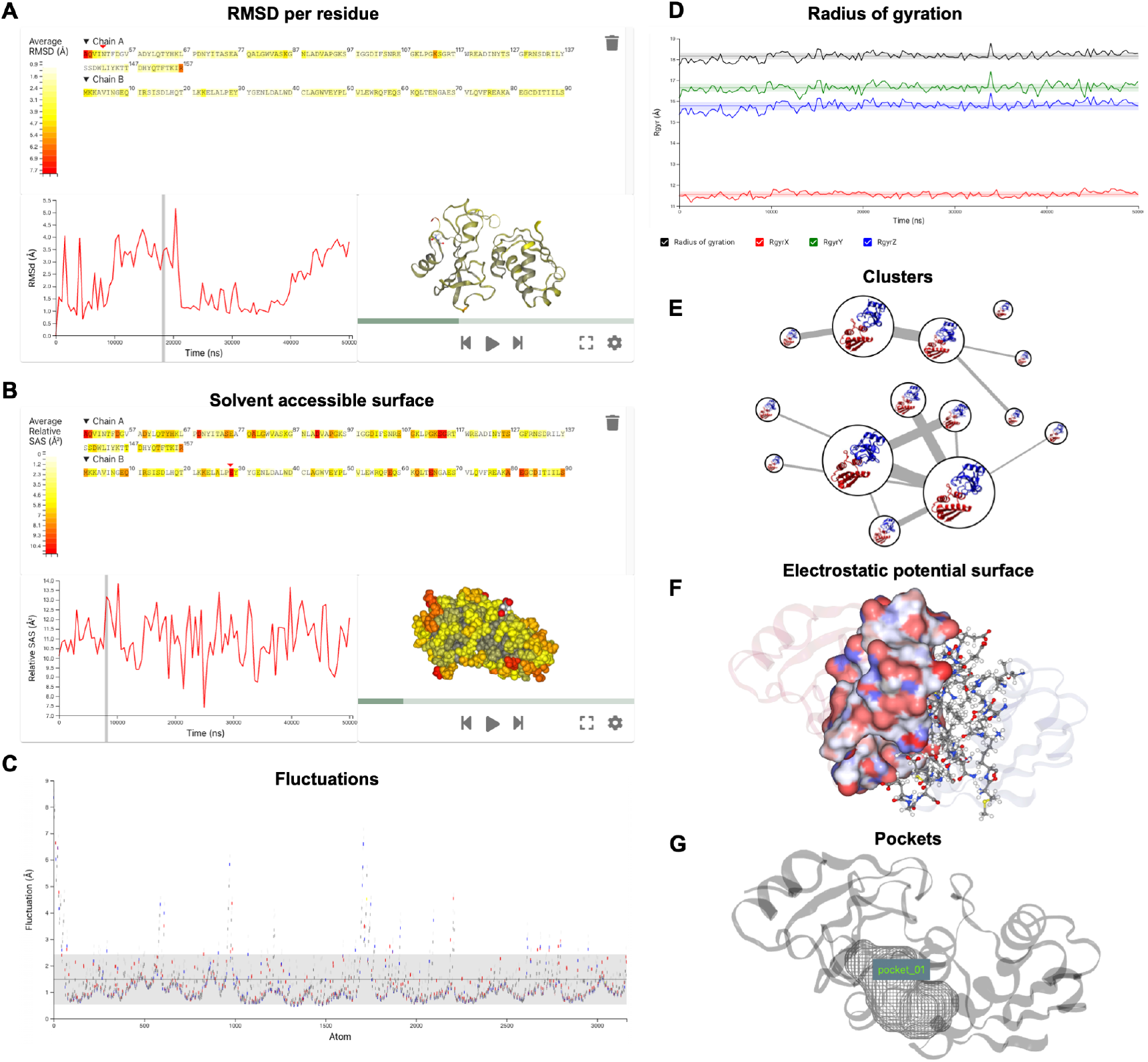
Example of pre-calculated analyses for PDB code 1B2S from the PDBbind dataset. The evolution of **A)** RMSD per residue, **B)** Solvent accessible surface, **C)** fluctuations, and **D)** radius of gyration are shown along the simulation time for the replica 1 of 1B2S simulation. **E)** The cluster analysis is shown here, where the size of the circle represents cluster population ratio. **F)** The electrostatic potential surface highlights the interacting residues. **G** Pockets are identified and mapped on the structure.

### Downloads page

Users can download three files for each entry including main structure (.pdb), main trajectory (.xtc), and main topology (.tpr) files. This page is also accessible through programmatic API, which is applicable specifically for large MD simulations.

## DISCUSSION

Recent breakthroughs in artificial intelligence have made it possible to extract rich structural information from macromolecules. However, a persistent and critical limitation of most models is their reliance on static representations, which fail to capture the dynamic nature of macromolecular function. This limitation is driven largely by the scarcity of high-quality, large-scale dynamic datasets. For example, state-of-the-art binding site predictors [45, 46] are trained on hundreds of thousands of static protein structures, yet no equivalent-scale dynamic resource exists. To address this gap, we joined the collective European MDDB initiative to generate and share FAIR-compliant MD data for macromolecular assemblies. By characterizing their conformational dynamics, DynaRepo bridges the gap between static structure and biological function.

DynaRepo provides all-atom MD trajectories for a large and diverse set of macromolecular complexes as well as single-chain proteins. The current release includes trajectories for more than 700 systems and 1,500 protein chains, totaling 1,146 *µ*s of simulation time. The systems were selected from PDBbind, antigens from the SAbDab database, the “Testing Transient” dataset, and structural variants of nucleosome complexes. DynaRepo is freely accessible at https://dynarepo.inria.fr/. Each entry is accompanied by pre-computed post-simulation analyses, including RMSD, RMSF, TM-score, radius of gyration, PCA, solvent-accessible surface, residue-wise distances, electrostatic potential surfaces, hydrogen bonds, energies, and pocket detection, available through interactive visualizations. As the first MDDB node in France, DynaRepo offers open access to data, analysis protocols, and simulation inputs, in full alignment with FAIR data principles.

The primary goal of DynaRepo is to enable the study of macromolecular dynamics at scale. We have already used these data to investigate key biological questions, including mapping communication networks and allosteric signaling [47]. We have also introduced a dynamic-aware deep learning architecture that combines cooperative graph neural networks [48] with a geometric transformer to predict binding sites in flexible regions [49], with benchmarks showing clear improvements when dynamics are incorporated. Beyond binding site prediction, the data in DynaRepo can be used to encode conformational flexibility for function annotation, interface detection, and binding affinity estimation, tasks that are essential for biology, biophysics, and therapeutic discovery. However, significant bottlenecks remain, most notably the shortage of dynamic datasets for protein–nucleic acid complexes, which currently limits broad generalization across molecular interaction types.

## Data availability

The database website is freely available online without login requirement at https://dynarepo.inria.fr/.

## ACKNOWLEDGEMENTS

This work was granted access to the HPC resources of IDRIS under the allocations 2024-GC010715460, 2023-A0150714577 and 2022-A0120713412 made by GENCI, and of the EXPLOR center (grant 2019CP-MXX0983). We are grateful to Adam Hospital, Daniel Beltran, and Genis Bayarri (IRB Barcelona) for their support in the database implementation, and providing the MDDB workflow, and to Frederic Beck (Service d’expérimentation et de développement (SED) at Centre Inria de l’Université de Lorraine) for support in running the post-simulation analyses. HK and EB were supported by the French Agence Nationale de la Recherche (ANR), under grants ANR-22-CPJ2-0075-01, ANR-24-CE45-4243-01, and ANR-24-CE29-2473. YK was supported by the French National Research Agency (ANR) under the France 2030 grant reference number ANR-24-RRII-0002 operated by the Inria Quadrant Program.

## Author Contributions

OM: Data curation, Formal analysis, Investigation, writing - original draft; EB: Data curation, Formal analysis; HK: Conceptualization, Formal analysis, Writing - original draft, Resources; YK: Conceptualization, Data curation, Formal analysis, Supervision, Writing - original draft.

## References

1. John Jumper, Richard Evans, Alexander Pritzel, Tim Green, Michael Figurnov, Olaf Ronneberger, Kathryn Tunyasuvunakool, Russ Bates, Augustin Žídek, Anna Potapenko, et al. Highly accurate protein structure prediction with alphafold. nature, 596(7873): 583–589, 2021.

2. Josh Abramson, Jonas Adler, Jack Dunger, Richard Evans, Tim Green, Alexander Pritzel, Olaf Ronneberger, Lindsay Willmore, Andrew J Ballard, Joshua Bambrick, et al. Accurate structure prediction of biomolecular interactions with alphafold 3. Nature, 630(8016): 493–500, 2024.

3. Minkyung Baek, Ryan McHugh, Ivan Anishchenko, Hanlun Jiang, David Baker, and Frank DiMaio. Accurate prediction of protein–nucleic acid complexes using rosettafoldna. Nature methods, 21(1):117–121, 2024.

4. Amitava Roy, Ethan Ward, Illyoung Choi, Michele Cosi, Tony Edgin, Travis S Hughes, Md Shafayet Islam, Asif M Khan, Aakash Kolekar, Mariah Rayl, et al. Mdrepo—an open data warehouse for community-contributed molecular dynamics simulations of proteins. Nucleic Acids Research, 53(D1):D477–D486, 2025.

5. Phillip J Stansfeld, Joseph E Goose, Martin Caffrey, Elisabeth P Carpenter, Joanne L Parker, Simon Newstead, and Mark SP Sansom. Memprotmd: automated insertion of membrane protein structures into explicit lipid membranes. Structure, 23(7): 1350–1361, 2015.

6. Ismael Rodríguez-Espigares, Mariona Torrens-Fontanals, Johanna KS Tiemann, David Aranda-García, Juan Manuel Ramírez-Anguita, Tomasz Maciej Stepniewski, Nathalie Worp, Alejandro Varela-Rial, Adrián Morales-Pastor, Brian Medel-Lacruz, et al. Gpcrmd uncovers the dynamics of the 3d-gpcrome. Nature Methods, 17(8): 777–787, 2020.

7. Mariona Torrens-Fontanals, Alejandro Peralta-García, Carmine Talarico, Ramon Guixa-González, Toni Giorgino, and Jana Selent. Scov2-md: a database for the dynamics of the sars-cov-2 proteome and variant impact predictions. Nucleic acids research, 50(D1):D858–D866, 2022.

8. Tim Meyer, Marco D’Abramo, Manuel Rueda, Carles Ferrer-Costa, Alberto Pérez, Oliver Carrillo, Jordi Camps, Carles Fenollosa, Dmitry Repchevsky, Josep Lluis Gelpí, et al. Model (molecular dynamics extended library): a database of atomistic molecular dynamics trajectories. Structure, 18(11): 1399–1409, 2010.

9. Marc W van der Kamp, R Dustin Schaeffer, Amanda L Jonsson, Alexander D Scouras, Andrew M Simms, Rudesh D Toofanny, Noah C Benson, Peter C Anderson, Eric D Merkley, Steven Rysavy, et al. Dynameomics: a comprehensive database of protein dynamics. Structure, 18(4): 423–435, 2010.

10. Yann Vander Meersche, Gabriel Cretin, Aria Gheeraert, Jean-Christophe Gelly, and Tatiana Galochkina. Atlas: protein flexibility description from atomistic molecular dynamics simulations. Nucleic acids research, 52(D1):D384–D392, 2024.

11. Zhihai Liu, Minyi Su, Li Han, Jie Liu, Qifan Yang, Yan Li, and Renxiao Wang. Forging the basis for developing protein-ligand interaction scoring functions. Acc. Chem. Res., 50(2):302–309, February 2017.

12. James Dunbar, Konrad Krawczyk, Jinwoo Leem, Terry Baker, Angelika Fuchs, Guy Georges, Jiye Shi, and Charlotte M Deane. Sabdab: the structural antibody database. Nucleic acids research, 42(D1):D1140–D1146, 2014.

13. João P. G. L. M. Rodrigues, João M.C. Teixeira, Mikaël Trellet, and Alexandre M. J. J. Bonvin. pdb-tools: a swiss army knife for molecular structures. F1000Research, 7:1961, December 2018.

14. Srayanta Mukherjee and Yang Zhang. Mm-align: a quick algorithm for aligning multiple-chain protein complex structures using iterative dynamic programming. Nucleic Acids Research, 37(11):e83–e83, May 2009.

15. P. Gainza, F. Sverrisson, F. Monti, E. Rodolà, D. Boscaini, M. M. Bronstein, and B. E. Correia. Deciphering interaction fingerprints from protein molecular surfaces using geometric deep learning. Nature Methods, 17(2):184–192, December 2019.

16. Nan Zhao, Bingqing Han, Cuicui Zhao, Jinbo Xu, and Xinqi Gong. Abag-docking benchmark: a non-redundant structure benchmark dataset for antibody–antigen computational docking. Briefings in Bioinformatics, 25(2), January 2024.

17. Martin Steinegger and Johannes Söding. Mmseqs2 enables sensitive protein sequence searching for the analysis of massive data sets. Nature Biotechnology, 35(11):1026–1028, October 2017.

18. Yasaman Karami and Emmanuelle Bignon. Cysteine hyperoxidation rewires communication pathways in the nucleosome and destabilizes the dyad. Computational and Structural Biotechnology Journal, 23: 1387–1396, 2024.

19. Ekaterina Smirnova, Emmanuelle Bignon, Patrick Schultz, Gabor Papai, and Adam Ben Shem. Binding to nucleosome poises human sirt6 for histone h3 deacetylation. Elife, 12:RP87989, 2024.

20. Yasaman Karami, Roy González-Alemán, Mailys Duch, Yuya Qiu, Yani Kedjar, and Emmanuelle Bignon. Histone h3 as a redox switch in the nucleosome core particle: insights from molecular modeling. bioRxiv, pages 2024–10, 2024.

21. David Van Der Spoel, Erik Lindahl, Berk Hess, Gerrit Groenhof, Alan E Mark, and Herman JC Berendsen. Gromacs: fast, flexible, and free. Journal of computational chemistry, 26(16): 1701–1718, 2005.

22. J Michael Word, Simon C Lovell, Jane S Richardson, and David C Richardson. Asparagine and glutamine: using hydrogen atom contacts in the choice of side-chain amide orientation. Journal of molecular biology, 285(4): 1735–1747, 1999.

23. Herman JC Berendsen, JPM van Postma, Wilfred F Van Gunsteren, ARHJ DiNola, and Jan R Haak. Molecular dynamics with coupling to an external bath. The Journal of chemical physics, 81(8):3684–3690, 1984.

24. Giovanni Bussi, Davide Donadio, and Michele Parrinello. Canonical sampling through velocity rescaling. The Journal of chemical physics, 126(1), 2007.

25. Mattia Bernetti and Giovanni Bussi. Pressure control using stochastic cell rescaling. The Journal of Chemical Physics, 153(11), 2020.

26. Berk Hess, Henk Bekker, Herman JC Berendsen, and Johannes GEM Fraaije. Lincs: A linear constraint solver for molecular simulations. Journal of computational chemistry, 18(12): 1463–1472, 1997.

27. Darrin M York, Tom A Darden, and Lee G Pedersen. The effect of long-range electrostatic interactions in simulations of macromolecular crystals: A comparison of the ewald and truncated list methods. The Journal of chemical physics, 99(10): 8345–8348, 1993.

28. D.A. Case, K. Belfon, I.Y. Ben-Shalom, S.R. Brozell, D.S. Cerutti, T.E. Cheatham III, V.W.D Cruzeiro, T.A. Darden, R.E. Duke, G. Giambasu, M.K. Gilson, H. Gohlke, A.W. Goetz, R. Harris, S. Izadi, S.A. Izmailov, K. Kasavajhala, A. Kovalenko, R. Krasny, T. Kurtzman, T.S. Lee, S. LeGrand, P. Li, C. Lin, J. Liu, T. Luchko, R. Luo, V. Man, K.M. Merz, Y. Miao, O. Mikhailovskii, G. Monard, H. Nguyen, A. Onufriev, F. Pan, S. Pantano, R. Qi, D.R. Roe, A. Roitberg, C. Sagui, S. Schott-Verdugo, J. Shen, C.L. Simmerling, N.R. Skrynnikov, J. Smith, J. Swails, R.C. Walker, J. Wang, L. Wilson, R.M. Wolf, X. Wu, Y. Xiong, Y. Xue, D.M. York, and P.A. Kollman. Amber 2020. 2020. University of California, San Francisco.

29. James C Phillips, David J Hardy, Julio DC Maia, John E Stone, João V Ribeiro, Rafael C Bernardi, Ronak Buch, Giacomo Fiorin, Jérôme Hénin, Wei Jiang, et al. Scalable molecular dynamics on cpu and gpu architectures with namd. J. Chem. Phys., 153(4): 044130, 2020.

30. James A Maier, Carmenza Martinez, Koushik Kasavajhala, Lauren Wickstrom, Kevin E Hauser, and Carlos Simmerling. ff14sb: improving the accuracy of protein side chain and backbone parameters from ff99sb. Journal of chemical theory and computation, 11(8): 3696–3713, 2015.

31. Ivan Ivani, Pablo D Dans, Agnes Noy, Alberto Perez, Ignacio Faustino, Adam Hospital, Jurgen Walther, Pau Andrio, Ramon Goni, Alexandra Balaceanu, Guillem Portella, Federica Battistini, Josep Lluis Gelpí, Carlos González, Michele Vendruscolo, Charles A Laughton, Sarah A Harris, David A Case, and Modesto Orozco. Parmbsc1: a refined force field for DNA simulations. Nature Methods, 38(13): 55–58, 2016.

32. Jejoong Yoo and Aleksei Aksimentiev. New tricks for old dogs: improving the accuracy of biomolecular force fields by pair-specific corrections to non-bonded interactions. Physical Chemistry Chemical Physics, 20(13): 8432–8449, 2018.

33. Pekka Mark and Lennart Nilsson. Structure and dynamics of the TIP3P, SPC, and SPC/E water models at 298 K. Journal of Physical Chemistry A, 105(43): 9954–9960, 2001.

34. Robert Zwanzig. Nonlinear generalized langevin equations. Journal of Statistical Physics, 9(3):215–220, 1973.

35. Scott E Feller, Yuhong Zhang, Richard W Pastor, and Bernard R Brooks. Constant pressure molecular dynamics simulation: The langevin piston method. The Journal of chemical physics, 103(11):4613–4621, 1995.

36. Shuichi Miyamoto and Peter A. Kollman. Settle: An analytical version of the SHAKE and RATTLE algorithm for rigid water models. Journal of Computational Chemistry, 13(8):952–962, oct 1992.

37. Chad W. Hopkins, Scott Le Grand, Ross C. Walker, and Adrian E. Roitberg. Long-time-step molecular dynamics through hydrogen mass repartitioning. Journal of Chemical Theory and Computation, 11(4):1864–1874, apr 2015.

38. Daniel Beltrán, Adam Hospital, Josep Lluís Gelpí, and Modesto Orozco. A new paradigm for molecular dynamics databases: the covid-19 database, the legacy of a titanic community effort. Nucleic Acids Research, 52(D1):D393–D403, 2024.

39. Yang Zhang and Jeffrey Skolnick. Scoring function for automated assessment of protein structure template quality. Proteins, 57(4):702–710, December 2004.

40. J L Gelpí, S G Kalko, X Barril, J Cirera, X de La Cruz, F J Luque, and M Orozco. Classical molecular interaction potentials: improved setup procedure in molecular dynamics simulations of proteins. Proteins, 45(4):428–437, December 2001.

41. Peter Schmidtke, Axel Bidon-Chanal, F Javier Luque, and Xavier Barril. MDpocket: open-source cavity detection and characterization on molecular dynamics trajectories. Bioinformatics, 27(23):3276– 3285, December 2011.

42. Rommie E Amaro, Johan Åqvist, Ivet Bahar, Federica Battistini, Adam Bellaiche, Daniel Beltran, Philip C Biggin, Massimiliano Bonomi, Gregory R Bowman, Richard A Bryce, et al. The need to implement fair principles in biomolecular simulations. Nature methods, pages 1–5, 2025.

43. Sarah Hunter, Rolf Apweiler, Teresa K Attwood, Amos Bairoch, Alex Bateman, David Binns, Peer Bork, Ujjwal Das, Louise Daugherty, Lauranne Duquenne, et al. Interpro: the integrative protein signature database. Nucleic acids research, 37(suppl 1):D211–D215, 2009.

44. Jonathan Lees, Corin Yeats, James Perkins, Ian Sillitoe, Robert Rentzsch, Benoit H Dessailly, and Christine Orengo. Gene3d: a domain-based resource for comparative genomics, functional annotation and protein network analysis. Nucleic acids research, 40(D1):D465–D471, 2012.

45. Pablo Gainza, Freyr Sverrisson, Frederico Monti, Emanuele Rodola, Davide Boscaini, Michael M Bronstein, and Bruno E Correia. Deciphering interaction fingerprints from protein molecular surfaces using geometric deep learning. Nature methods, 17(2): 184–192, 2020.

46. Lucien F Krapp, Luciano A Abriata, Fabio Cortés Rodriguez, and Matteo Dal Peraro. Pesto: parameter-free geometric deep learning for accurate prediction of protein binding interfaces. Nature communications, 14(1): 2175, 2023.

47. Sneha Bheemireddy, Roy González-Alemán, Emmanuelle Bignon, and Yasaman Karami. Communication pathway analysis within protein-nucleic acid complexes. Journal of Chemical Theory and Computation, 2025.

48. Ben Finkelshtein, Xingyue Huang, Michael Bronstein, and Ismail Ilkan Ceylan. Cooperative graph neural networks. arXiv preprint arXiv:2310.01267, 2023.

49. Omid Mokhtari, Sergei Grudinin, Yasaman Karami, and Hamed Khakzad. Dynamicgt: a dynamic-aware geometric transformer model to predict protein binding interfaces in flexible and disordered regions. bioRxiv, pages 2025–03, 2025.

