## Supplementary Information for "DynaRepo: The repository of macromolecular conformational dynamics"

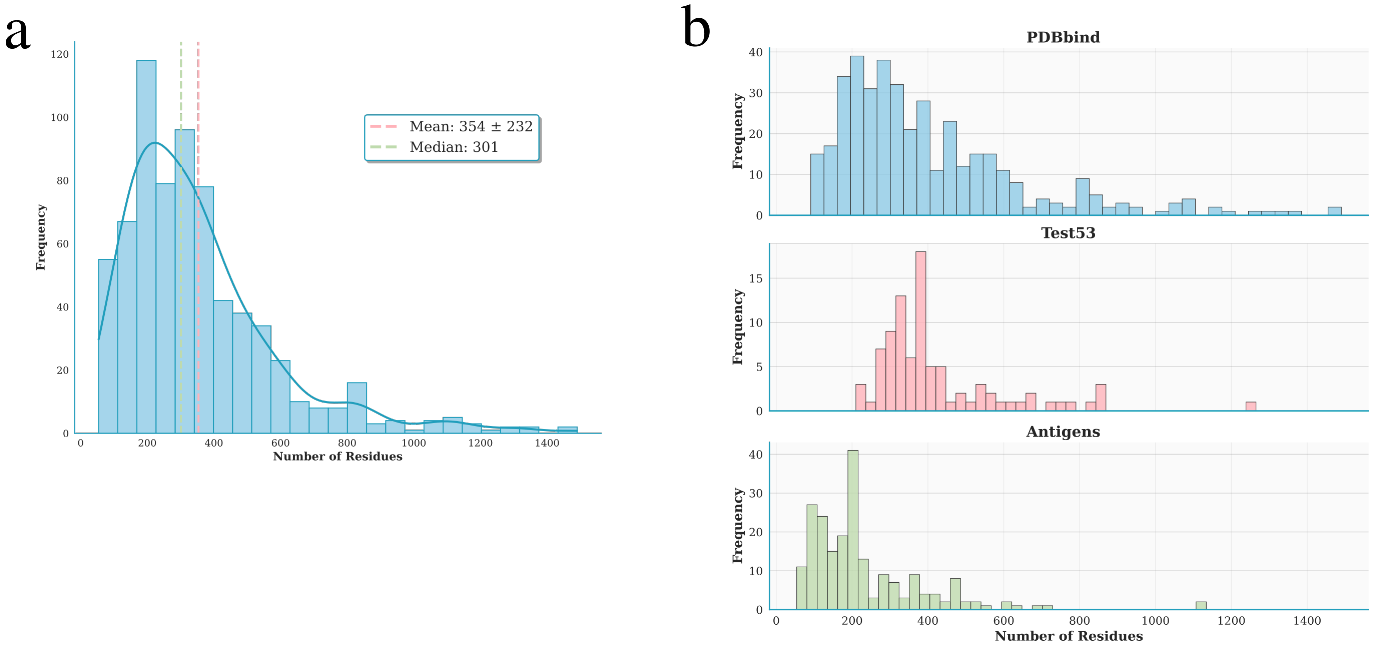


Figure S1: **Distribution of entry sizes.** (a) Overall distribution of the number of residues per MD simulation. (b) Same distribution shown separately for the PDBbind, Test53, and Antigen datasets.

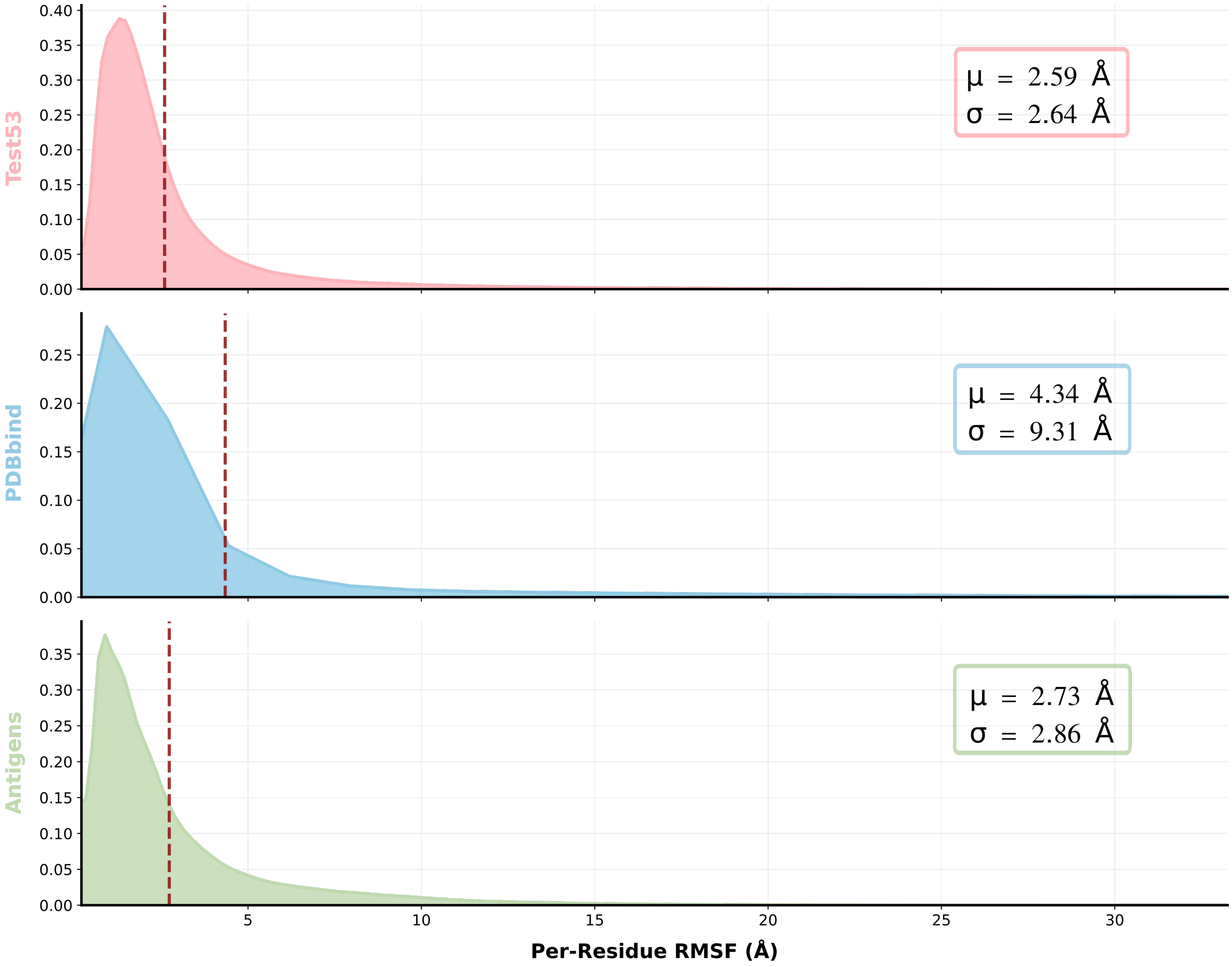


Figure S2: **Fluctuation analysis of dataset.** Probability density plot of per-residue fluctuations, with mean and standard deviation indicated. Extreme outliers were removed by excluding the top and bottom 1 percentile.

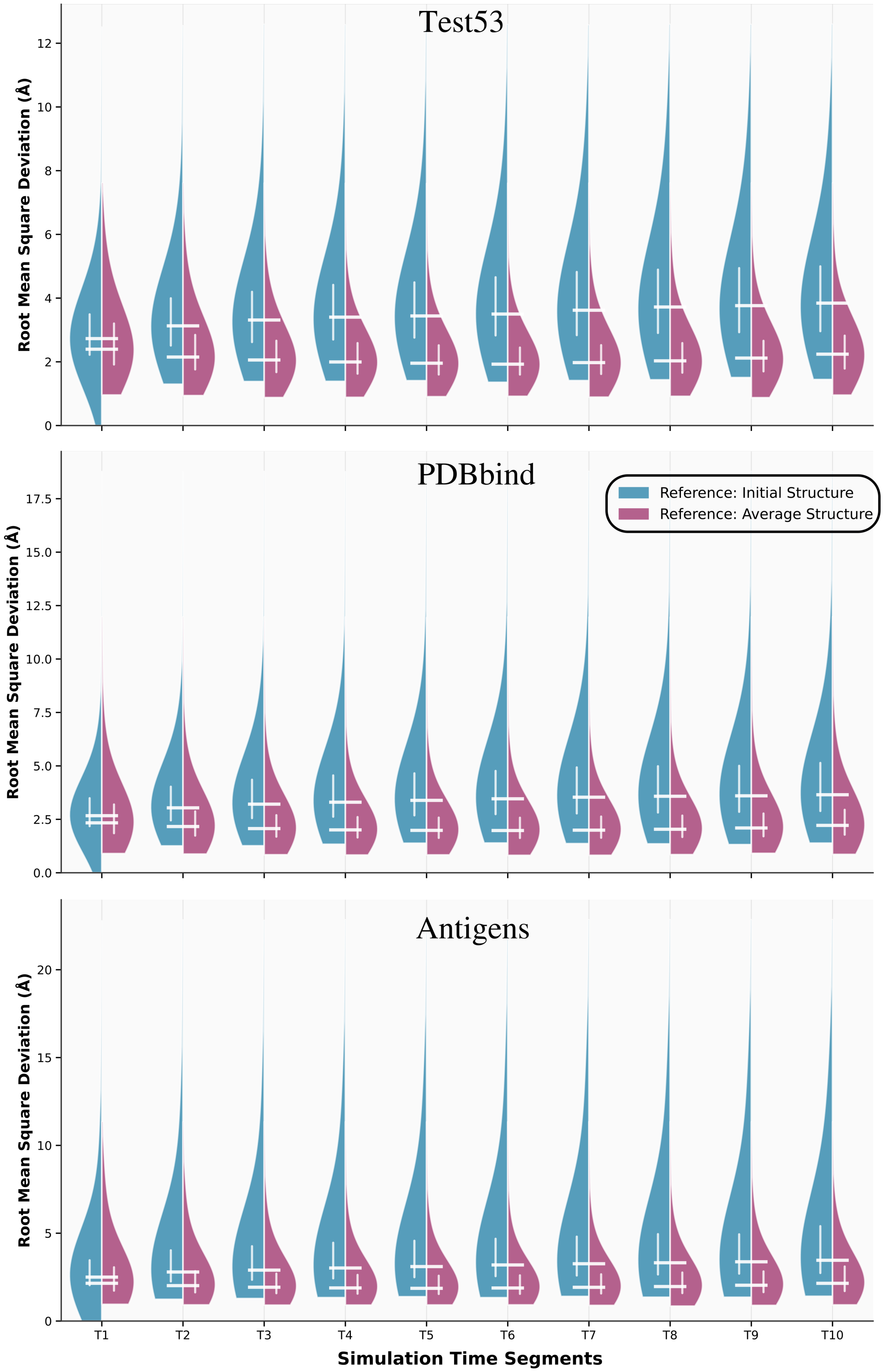


Figure S3: **RMSD analysis of dataset.** Violinplot showing the RMSD temporal variation across molecular dynamics simulations of dataset. Outliers are removed and frames from each replicate were analyzed independently.

### Illustrative Examples of DynaRepo Entries and Analyses

Here, we illustrate two example entries to showcase the different components of a DynaRepo dataset: **(i)** "COMPLEX OF THE HUMAN MHC CLASS I GLYCOPROTEIN HLA-A2 AND THE T CELL CORECEPTOR CD8" (PDB ID: 1AKJ), and **(ii)** "Nucleosome Core Particle, NCP147" (PDB ID: 1KX5).

#### Overview Page

The overview page provides general metadata associated with each entry. For example, users can: Select and view individual replicas (Fig. S3.1) which can later be used to explore the analysis specific to each replica, access the original PDB structure linked to the MD simulation (Fig. S3.2), inspect the length of the simulation (Fig. S3.3), view information about the submitting authors and the MD software used (Fig. S3.4).

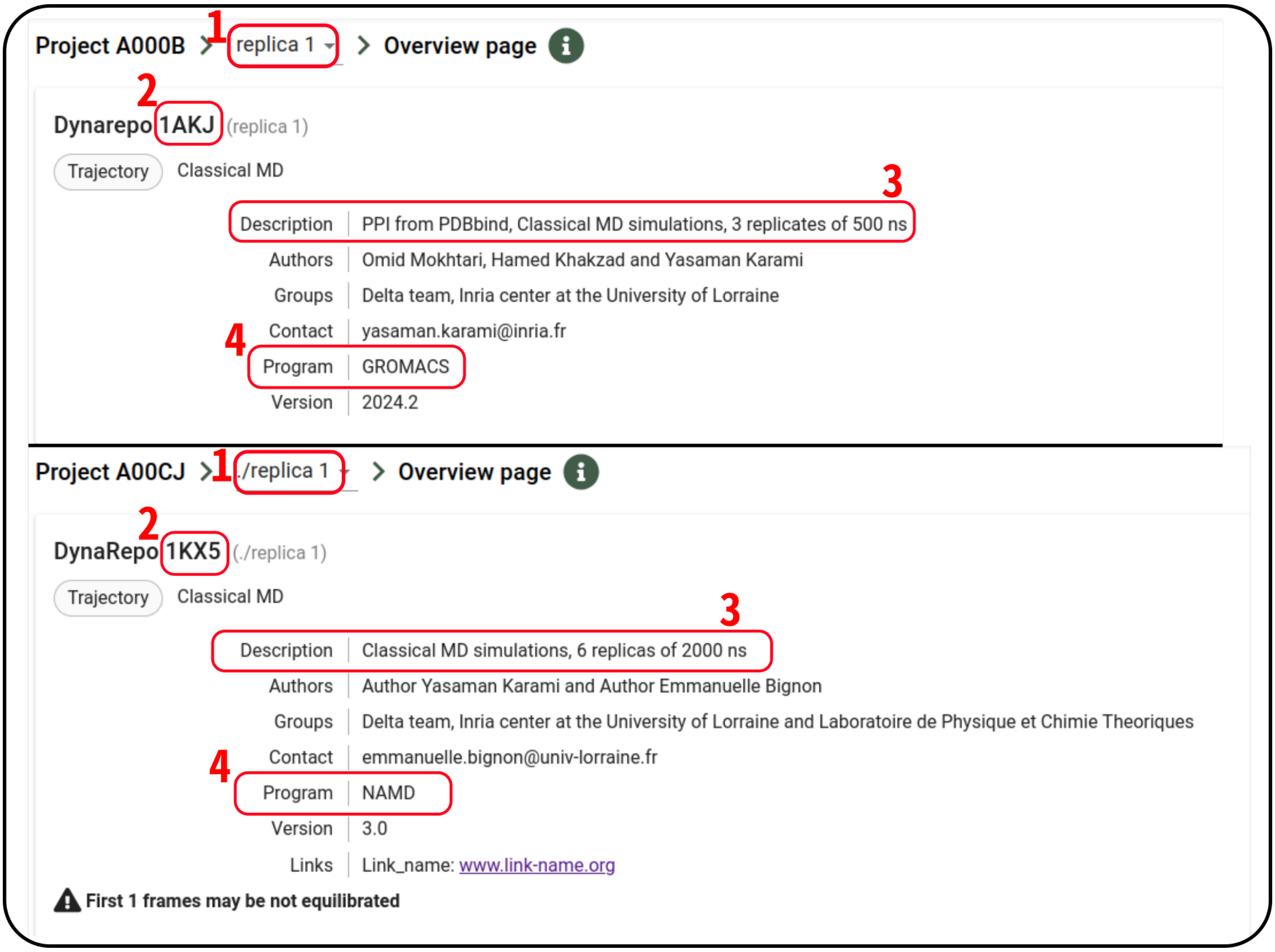


Figure S4: **Overview page.** Example entries for PDB IDs 1AKJ and 1KX5 showing general metadata including simulation length, replica selection, and MD software.

### Trajectory Page

The trajectory page enables in-browser visualization of the MD simulation without requiring download. Users can: **(i)** Filter by specific domains or chains (Fig. S4.1), and **(ii)** Customize visual settings such as coloring, rendering styles, background and other advanced options (Fig. S4.2).

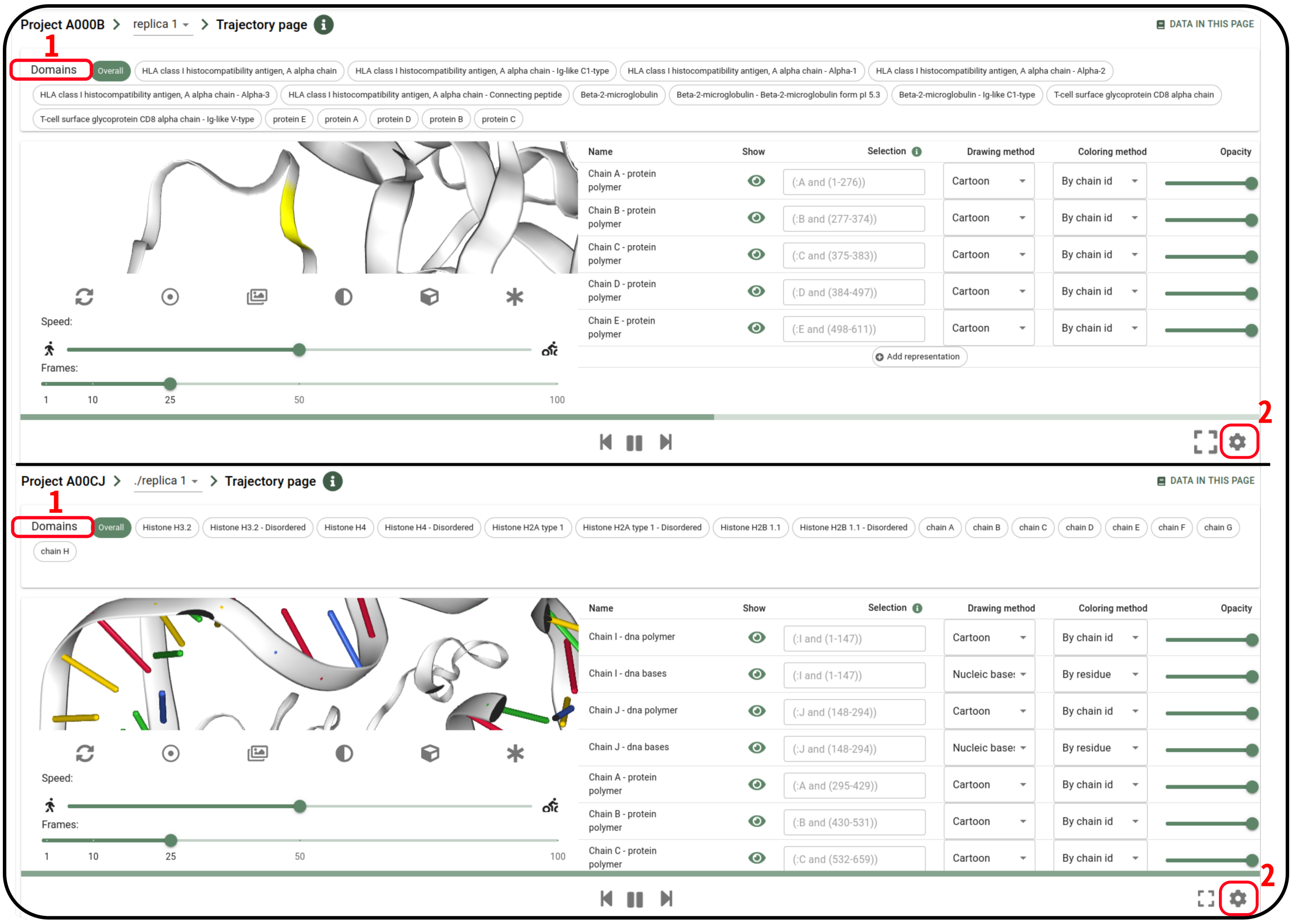


Figure S5: **Trajectory viewer.** Interactive visualization of the MD trajectory with options to filter by domain and customize display settings.

### Analysis Page

This section provides multiple categories of analyses, including: **(i)** Quality control, **(ii)** Interaction analyses, and **(iii)** Other.

In the *"RMSD per residue"* panel, users can explore residue-specific RMSD values across the simulation. Individual residues can be interactively selected to visualize their positional fluctuations over time (Fig. S5.1). The *"RMSDs"* section displays global structural deviations, including RMSD and TM-score values per frame relative to either the first frame or the average structure (Fig. S5.2).

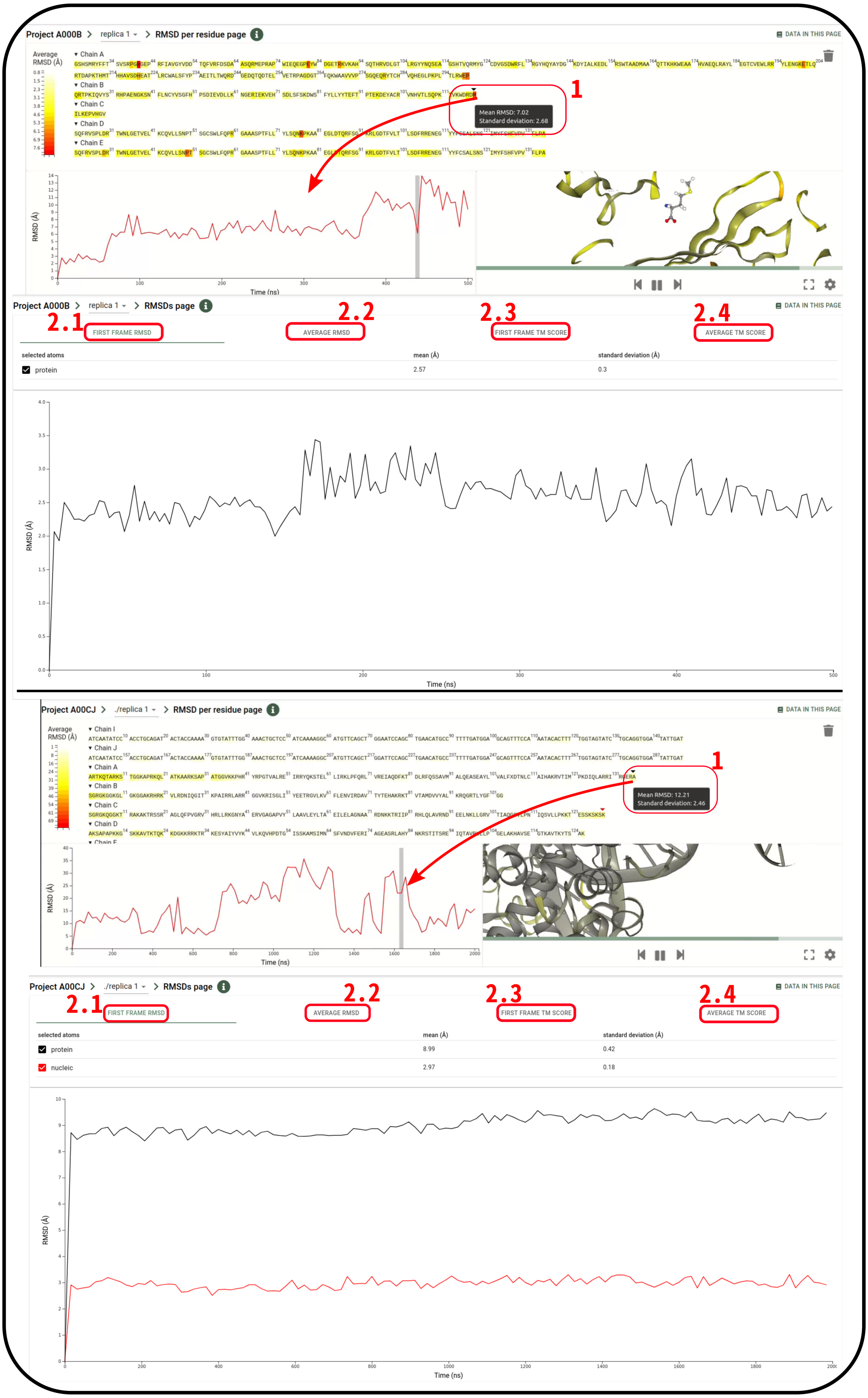


Figure S6: **RMSD analyses.** **Top:** Per-residue RMSD distribution with interactive residue tracking. **Bottom:** Frame-wise RMSD and TM-score plotted relative to the first and average frames.

The *"Radius of Gyration"* section displays how the compactness of the structure evolves over time. Line plots show the radius of gyration (Rg) across all simulation frames, with the mean and standard deviation highlighted. Users can identify frames with either significant Rg fluctuations (Fig. S6.1) or stable compactness (Fig. S6.2).

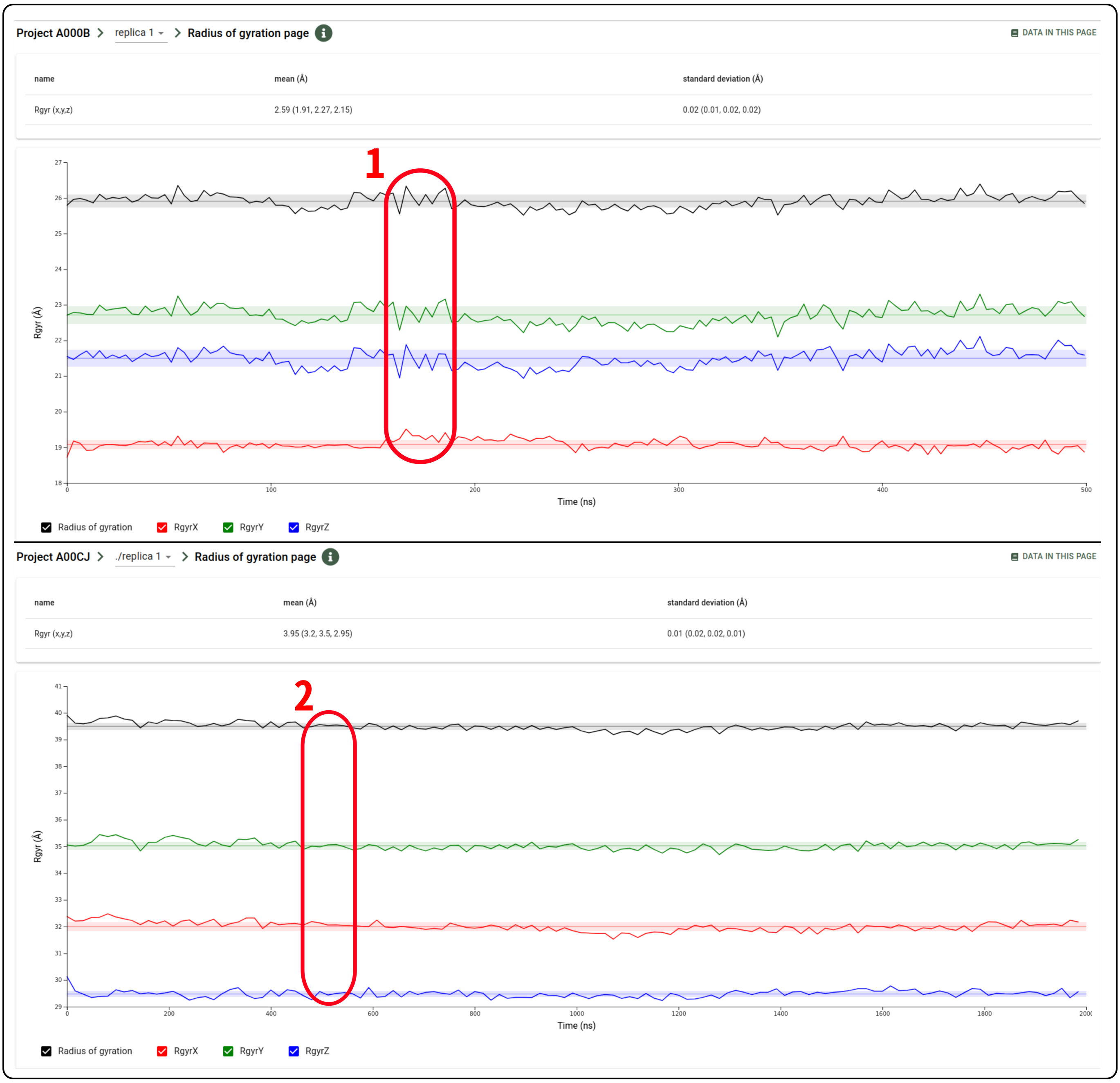


Figure S7: **Radius of Gyration analysis.** Rg values plotted over time with indication of average and variation across frames.

The *"Fluctuation"* page presents per-atom fluctuations across the simulation, with mean and standard deviation shown in Å. In the nucleosome example, clear differences in mobility between histone protein atoms and DNA atoms can be observed.

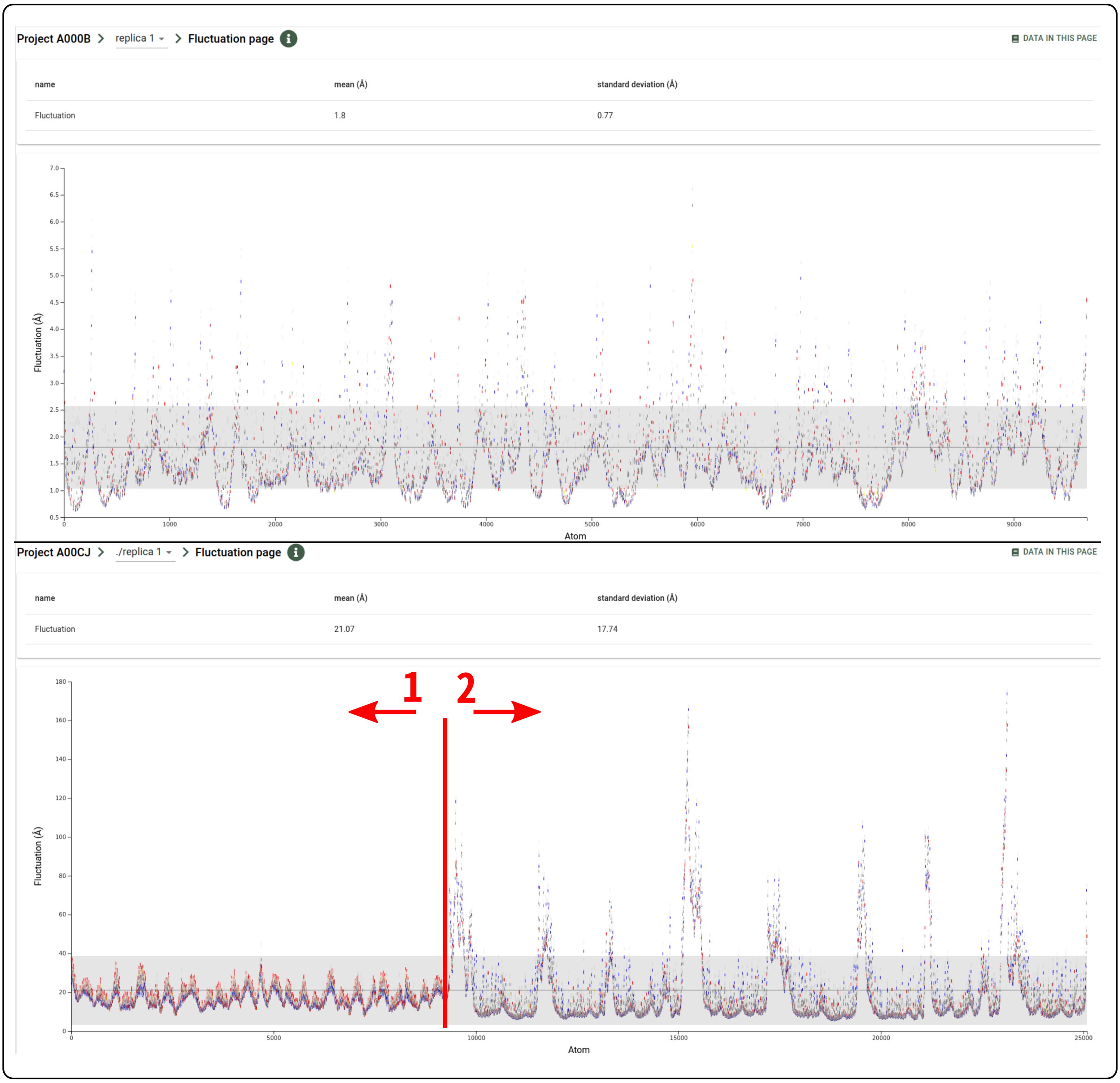


Figure S8: **Fluctuation analysis.** Atom-wise fluctuations with mean and standard deviation highlighted.

The PCA section displays a bar plot of eigenvalues corresponding to each principal component, representing the variance captured by the MD simulation. The yellow curve shows the cumulative explained variance across components.

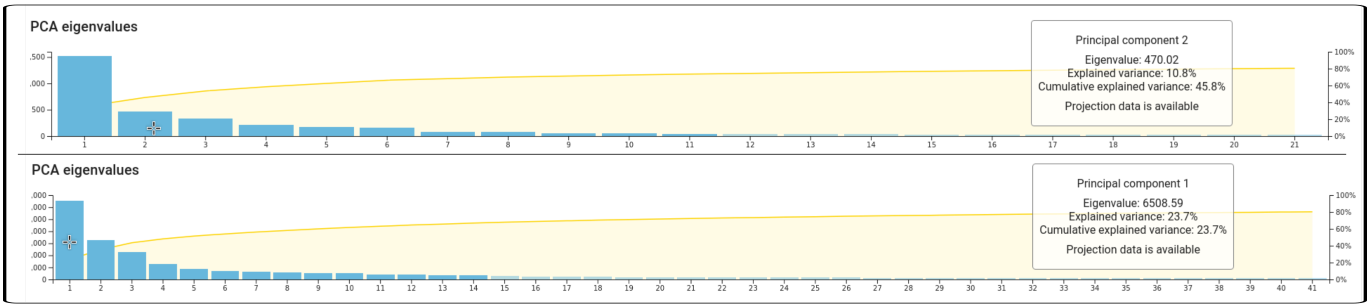


Figure S9: **PCA analysis.** Eigenvalues of principal components with cumulative variance (yellow curve).

The Solvent Accessible Surface (SAS) section presents the mean and variation of average SAS values per residue. Users can interactively select one or more residues (Fig. S9.1) to focus the visualization on these regions (Fig. S9.2) and observe SAS changes across frames via a line plot.

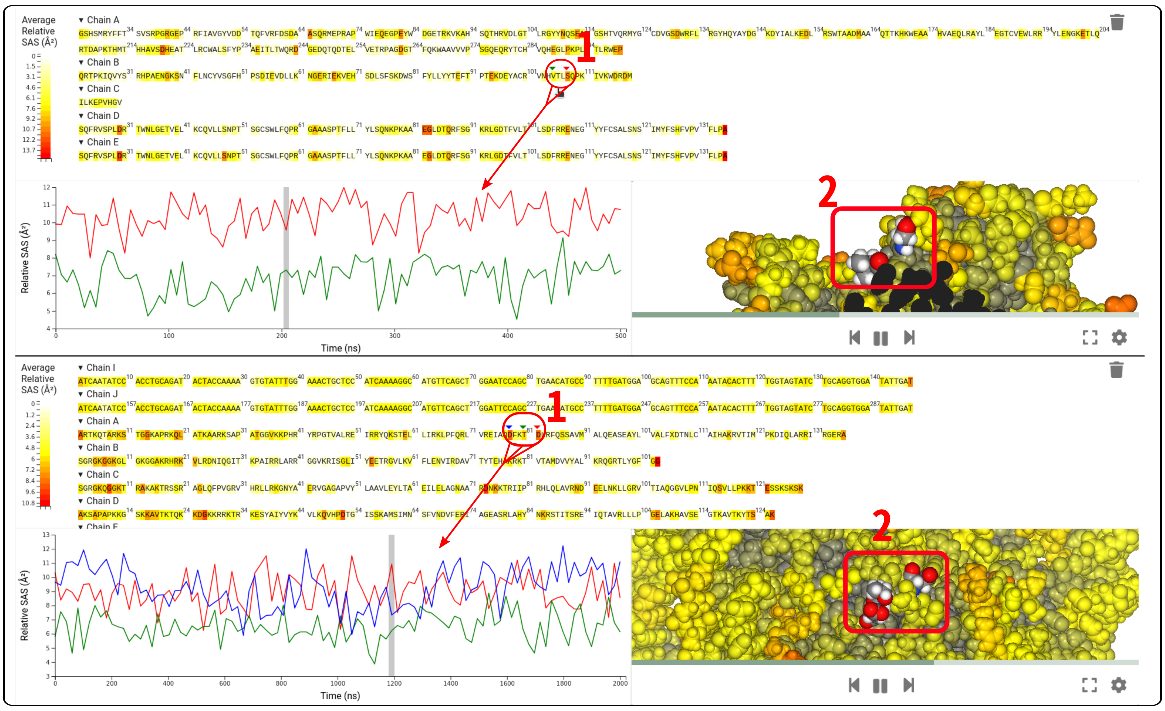


Figure S10: **Solvent accessible surface analysis.** Residue-wise mean and standard deviation of SAS values alongside interactive visualization.

The clustering section groups simulation frames based on structural similarity. Each node represents a representative frame from a cluster, with edges indicating structural proximity. The size of each node reflects the population of its corresponding conformational state. Users can interactively adjust the number of nodes or links to explore the structural landscape and transitions between states.

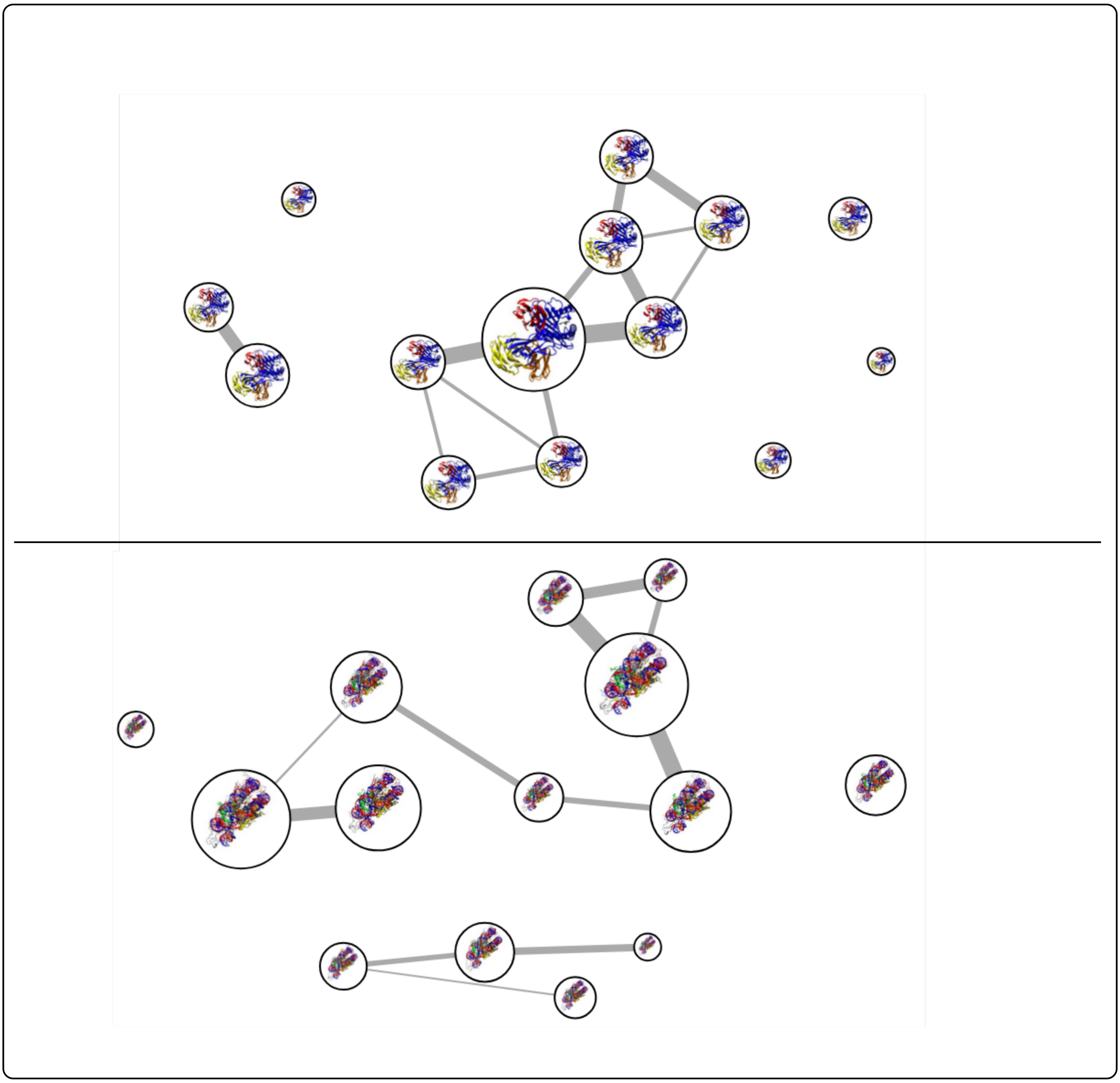

Figure S11: **Clustering analysis.** Representative frames grouped by structural similarity. Node size reflects cluster population; edge thickness indicates proximity between clusters.

*Distance per Residue* shows pairwise residue distances as heatmaps, reporting both average and standard deviation values. Users can select any pair of chains in the complex (e.g., protein–protein, protein–ligand) and optionally filter the map to show only interface residues (Fig. S11.1).

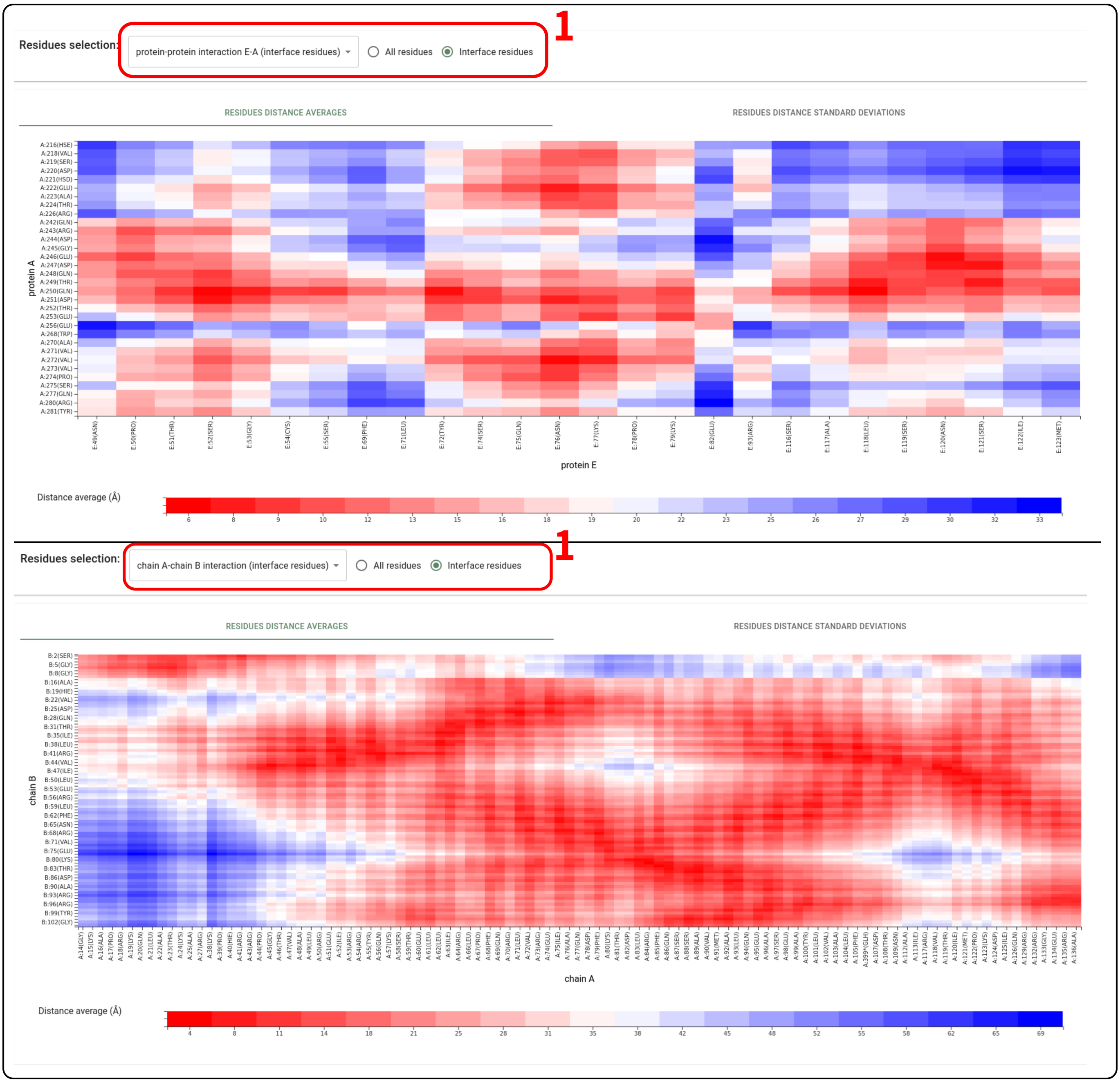


Figure S12: **Interaction analysis: Distance per residue.** Heatmap of average and standard deviation of pairwise residue distances between chains. Can be filtered to show only interface residues.

The *Electrostatic Potential Surface* section highlights interface residues between pairs of entities. One molecule is shown as a surface colored by atomic charges, while the interacting partner is rendered as balls and sticks to emphasize the interaction interface.

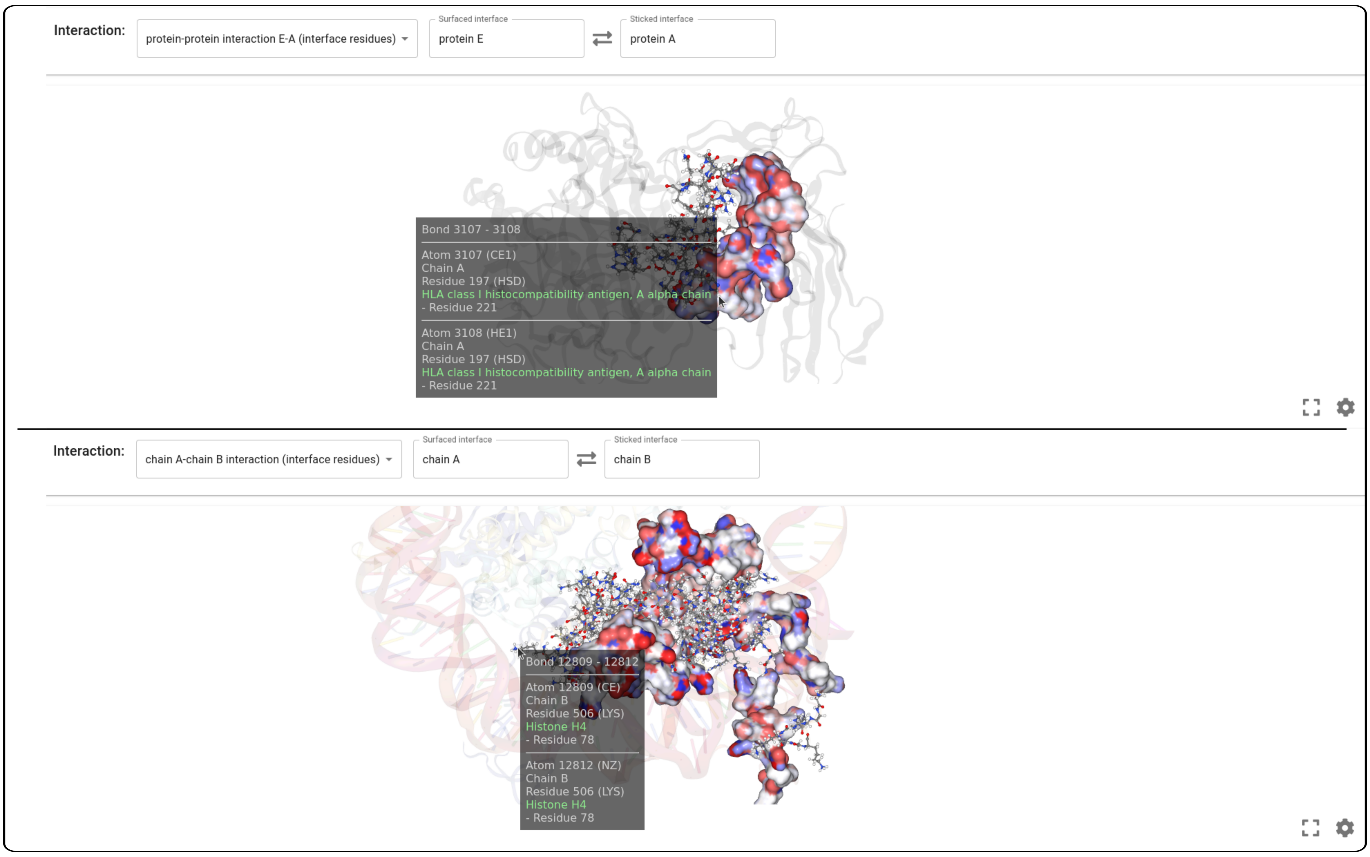


Figure S13: **Interaction analysis: Electrostatic potential surface.** Interface visualization with one partner shown as a surface and the other in balls-and-sticks representation.

48 In the *Hydrogen Bonds* section, all inter-entity hydrogen bonds are annotated. These interactions can be visualized directly on the structure, with distances between donor and acceptor atoms shown interactively.

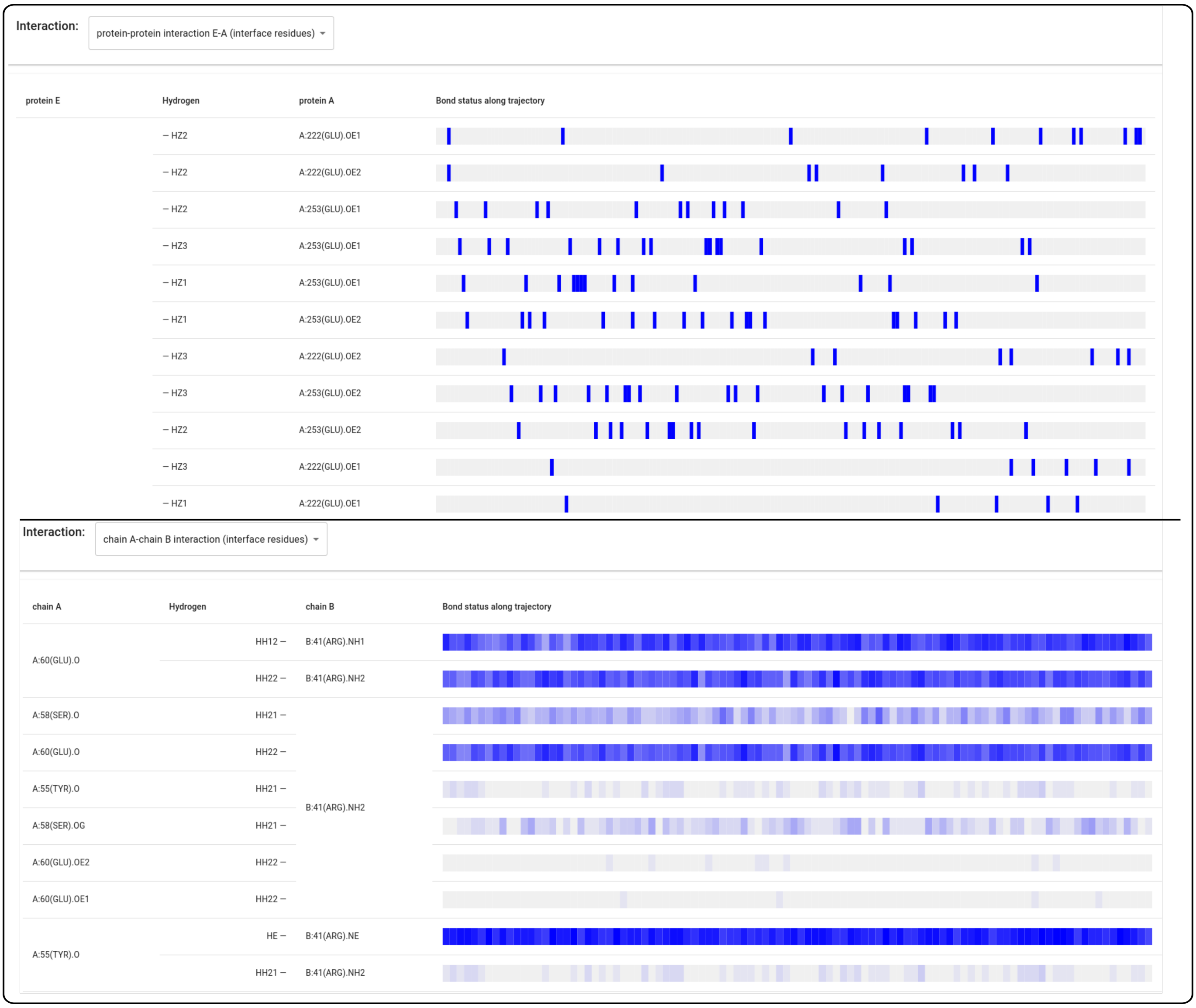


Figure S14: **Interaction analysis: Hydrogen bonds.** Annotated hydrogen bonds between chains shown with atomic distance visualization.

The *Pockets* panel displays detected pockets within the structure. Users can track pocket volume changes across the simulation via a line plot.

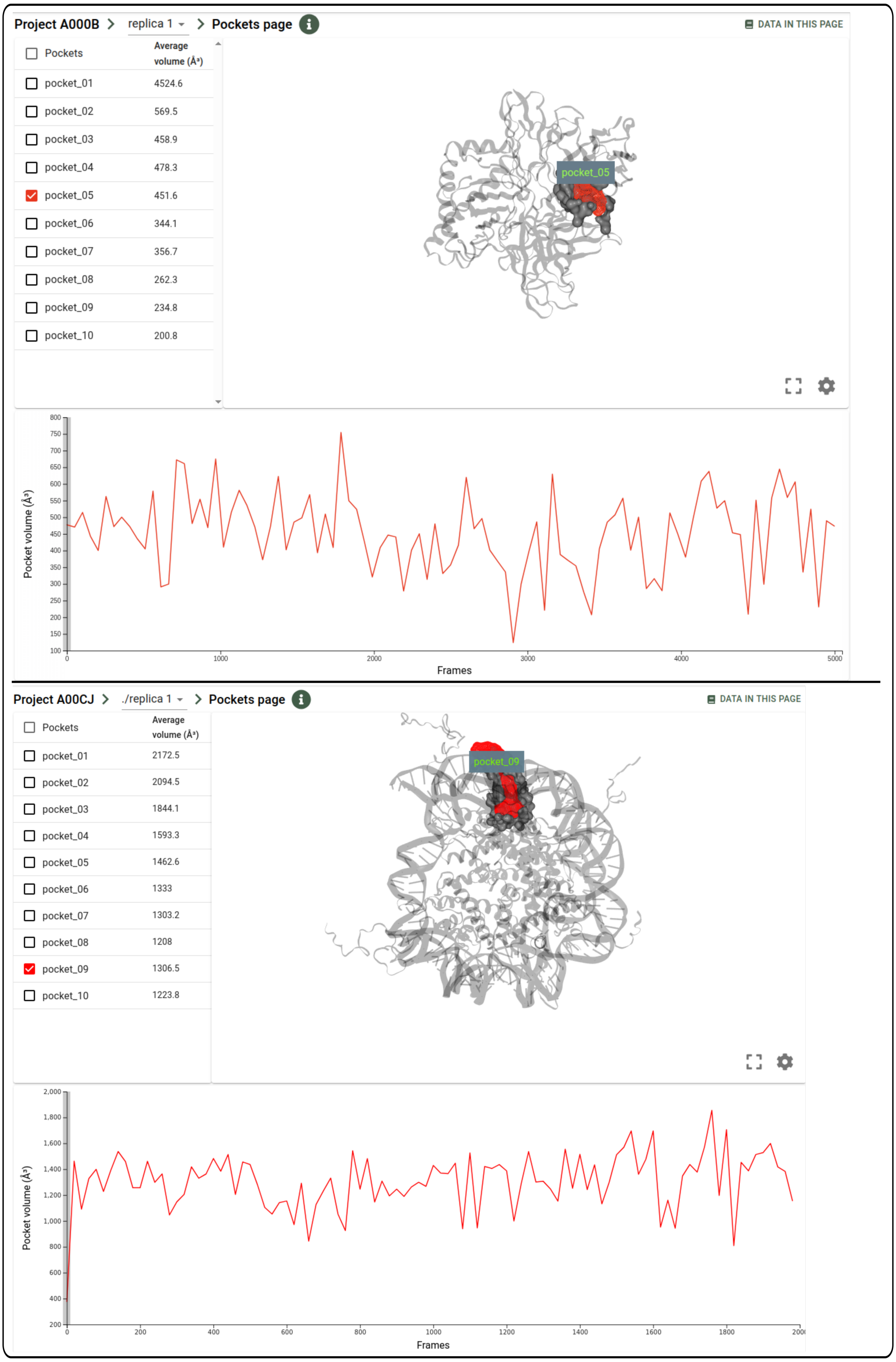


Figure S15: **Pocket analysis.** Detected pockets visualized with volume evolution plotted over time.
